# Comprehensive mapping of Cystic Fibrosis mutations to CFTR protein identifies mutation clusters and molecular docking predicts corrector binding site

**DOI:** 10.1101/242073

**Authors:** Steven V. Molinski, Vijay M. Shahani, Adithya S. Subramanian, Stephen S. MacKinnon, Geoffrey Woollard, Marcon Laforet, Onofrio Laselva, Leonard D. Morayniss, Christine E. Bear, Andreas Windemuth

## Abstract

**Background:** Cystic Fibrosis (CF) is caused by mutations in the *CFTR* gene, of which over 2000 have been reported to date. Mutations have yet to be analyzed in aggregate to assess their distribution across the tertiary structure of the CFTR protein, an approach that could provide valuable insights into the structure-function relationship of CFTR. In addition, the binding site of Class I correctors (VX-809, VX-661, C18) is not well understood.

**Methods:** Exonic *CFTR* mutations and mutant allele frequencies described in three curated databases (ABCMdb, CFTR1 and CFTR2, comprising >130,000 data points) were mapped to two different structural models: a homology model of full-length CFTR protein in the open-channel state, and a cryo-electron microscopy core-structure of CFTR in the closed-channel state. Immunoblotting confirmed the approximate binding site of Class I correctors, and molecular docking generated binding poses for their complex with the cryo-electron microscopy structure.

**Results:** Residue positions of six high-frequency mutant *CFTR* alleles were found to spatially co-localize in CFTR protein, and a significant cluster was identified at the NBD1:ICL4 interdomain interface. Further, Class I correctors VX-809, VX-661 and C18 were shown to act via a similar mechanism *in vitro*, and a putative multi-domain corrector binding site near residues F374-L375 was predicted *in silico*.

**Conclusions:** Our results confirm the significance of interdomain interfaces as susceptible to disruptive mutation, and identify a putative corrector binding site. The structural pharmacogenomics approach of mapping mutation databases to protein models shows promise for facilitating drug discovery and personalized medicine for monogenetic diseases.

## Introduction

Cystic Fibrosis (CF) is caused by mutations in the *cystic fibrosis transmembrane conductance regulator* (*CFTR/ABCC7*) gene which lead to defective biosynthesis, trafficking and/or activity of the CFTR protein^1,2^. CFTR is a chloride channel, comprised of five structural domains: membrane-spanning domain 1 (MSD1), nucleotide-binding domain 1 (NBD1), a regulatory (R) domain, MSD2 and NBD2. Together these domains cooperate to facilitate fluid transport and surface hydration across epithelial cells of the body’s tubular organs (e.g. lungs, intestines)^2,3^. Loss of CFTR expression and/or activity via mutation leads to a severe disease phenotype, where inflammation and infection within the lungs and intestines run rampant, and thus these organs are unable to sufficiently exchange gases and absorb nutrients, respectively^2–4^. To date, over 2000 disease-causing *CFTR* mutations have been reported, most of which are rare. The most common, F508del, is present in approximately 70% of the worldwide CF population (estimated to be 100,000); ~5% have G551D, ~5% G542X, and the remaining ~20% have one of the ~2000 other mutations^5–8^. These mutations are catalogued across three databases (ABCMdb^25^, abcmutations.hegelab.org; CFTR1, genet.sickkids.on.ca; CFTR2^7^, cftr2.org) and provide a comprehensive resource supporting the effort to further our understanding of the complex structure-function relationship of CFTR.

In recent years, much effort has been directed towards discovery and development of novel small molecule therapies for CF patients bearing the F508del and G551D mutations, since these mutations capture a large fraction of the patient population^9–13^. Accordingly, two small molecule therapies (Kalydeco®, comprised of the potentiator ivacaftor or VX-770, and Orkambi®, comprised of ivacaftor and the corrector lumacaftor or VX-809) have been developed for individuals with these mutations (G551D and F508del, respectively), as well as for those having a few other mutations with similar CFTR protein defects^14,15^. However, these discoveries as well as most other investigations typically explore *CFTR* mutations on a case-by-case basis, and do not consider the global mutation landscape as a whole. This is partly due to the abundance of the deposited data, which currently comprises a total count of >200,000 unique data points. Therefore, there is a need for studies which probe the landscape of known *CFTR* mutations in order to uncover salient trends and provide additional insight into the biochemical, biophysical and/or evolutionary nature of this fatal genetic disease.

Accordingly, we employed a structural pharmacogenomics approach^16,17^ to map all exonic mutations and mutant allele frequencies found in these databases of *CFTR* mutations onto three-dimensional structural models of the CFTR protein in both the open-channel^18^ and closed-channel^19^ states. The spatial distribution of the mapped mutations was analyzed to uncover salient trends that may inform on their mechanistic underpinnings. In addition, structural information from the experimentally determined closed-channel conformation^19^ was combined with computational docking calculations to predict CFTR residues that contribute to the binding sites of current small molecule therapeutics, mainly Class I correctors: VX-809 (lumacaftor, a component of Orkambi® co-therapy), VX-661 (tezacaftor, the putative successor of lumacaftor) and C18 (an investigational compound).

The proposed structural pharmacogenomics approach is shown to yield useful insights into structure-function relationships that underlie CF, and to predict a potential drug binding pocket which may facilitate further stratification of patients based on their mutation-sensitive ‘theratype’ or responsiveness to certain small molecule therapies^13,20^.

## Materials and Methods

### CFTR Mutation Databases

The three CFTR mutation databases used in this study are: ABCMdb^21^ (abcmutations.hegelab.org), CFTR1 (www.genet.sickkids.on.ca) and CFTR2^7^ (www.cftr2.org). Databases include mutations reported in the scientific literature (i.e. ABCMdb: containing validated disease-causing mutations, unvalidated disease-associated mutations, as well as experimental mutations used to study CFTR), those catalogued by physicians and scientists (i.e. CFTR1: validated and unvalidated mutations; CFTR2: validated mutations) as well as allele frequencies for reported mutations (CFTR2); databases were accessed between August 1^st^ and August 31^st^, 2016.

### CFTR Protein Tertiary Structures

We used a previously described and biochemically validated structural model of full-length CFTR in the open-channel state^18^ (i.e. residues 1–1480, including the extensively modeled R-domain), as well as the recent cryo-electron microscopy (cryo-EM)-derived partial/core-structure of CFTR at 3.87 Å resolution in the closed-channel state^19^ (lacking ~297 CFTR residues, including amino acids 1–4, 403–438, 646–843, 884–908, 1173–1206 and 1437–1480; PDB ID: 5UAK), as templates to map exonic *CFTR* mutations and allele frequencies. The homology model of CFTR in the open-channel state by Mornon *et al*. (2009)^18^ was used, rather than Mornon *et al*. (2015)^20^, as it is the only structural model of full-length CFTR (i.e. containing all 1480 residues); the latter model lacks the R-domain (defined as residues 650–843 in their study; 194 amino acids or ~13% of the protein) and subsequently would omit a substantial proportion of *CFTR* mutations described in the three CFTR mutation databases (i.e. ABCMdb, CFTR1, CFTR2). Further, use of a homology model was necessary since no experimental structure of CFTR in the open-channel state has been solved at sufficient resolution (i.e. the available low-resolution, ~9 Å structure^21^ does not have residue-level detail required for this study). In addition, although of lower resolution than desired for molecular docking approaches, the experimental cryo-EM structure of CFTR in the closed (inactive) conformation was used as a template for molecular docking studies, since it is well-established that ‘activating’ ligands (e.g. correctors) bind to *at least* this conformational state^5,24,25^.

### Mapping CFTR Mutations onto the Protein Tertiary Structures

CFTR mutations from the above listed databases were mapped onto the open- and closed-channel structures of CFTR. The ABCMdb, CFTR1 and CFTR2 mutation databases provided a detailed list of 1,955, 1,308 and 235 exonic mutations (missense and insertions/deletions, i.e. non-intronic), respectively, which allowed for mutation counts (i.e. number of unique missense or nonsense mutations at a given amino acid position) and allele frequencies to be tabulated and further highlighted on the CFTR protein model using an automated ‘paint-by-number’ approach. PyMOL (version 1.7) was used to visualize CFTR protein structures.

### Mutation Density using Average Interatomic Distance Measurements

To quantify the probability that the positions of the highest-frequency mutations (i.e. F508del, G542X, G551D, R553X W1282X, N1303K), known to be detrimental to structure and/or function, cluster within the CFTR tertiary structure, we computed the average interatomic distance (AID; mean value of distances between the C_α_ atoms of groups of amino acids) of residues F508, G542, G551, R553 W1282 and N1303, using the atomic coordinates of the full-length CFTR model of the active, open-channel state^18^. This distance was compared to the AID values of 1,000,000 randomly-selected six-residue subsets within the CFTR structural model to determine the statistical significance of the spatial clustering observed for the considered mutated positions.

### Identification of Interdomain Interface Residues using Proximity Measurements

PyMOL (version 1.7) was used to visualize interdomain interfaces in the full-length homology-based structural model of CFTR, mainly NBD1 (residues 381–630), NBD2 (residues 1171–1480) and MSD2 (i.e. intracellular loop 4 or ICL4, residues 1039–1093). Residues at domain-domain interfaces were defined as residues of one domain with at least one atom within a 4 Å distance of any atom in the neighboring domain. Interdomain interface residues for NBD1:NBD2 and NBD1:ICL4 were further annotated by mutation frequency, and residues contained within interdomain interfaces were coloured according to mutation counts.

### Molecular Docking of Pharmacological Correctors to CFTR

VX-809, VX-661 and C18 were docked to the apo form of human CFTR in the closed-channel state^19^ (PDB ID: 5UAK) using the CROSSx molecular docking tool from the Ligand Express™ computational suite (Cyclica Inc., Toronto, Canada)^26^. The lowest-energy conformers of the three correctors considered in isolation were modeled, and found to possess up to 20 Å of maximum possible length. Accordingly, a 30×30×30-Å cube centered around residues F374-L375 (amino acids shown to directly interact with the Class I corrector VX-809^27^) was used to define potential binding site region in the cryo-EM structure of CFTR. The small molecules were then docked to this region using a protocol, which allowed for flexibility of the ligand but kept the side-chains of target protein fixed (i.e. ‘rigid’ docking). A diverse ensemble of top-scoring sites centered around residues F374-L375 was visually inspected for chemical complementarity between the ligand and target moieties, and a reduced set of structures was docked allowing for flexible side-chains (i.e. ‘flexible’ docking). This protocol produced a total of 9 best docking poses for each of the corrector molecules. Assuming a common recognition mode for the three ligands, poses that align the pharmacophore features shared by the three corrector ligands, denoted here as the common scaffold (**Figure S4A**), were selected as the final docking solutions. PyMOL (version 1.7) was used to visualize molecular docking results.

### Quantifying CFTR MSD1 Protein Abundance

Human embryonic kidney (HEK)-293 cells were transfected with CFTR MSD1 constructs (i.e. full-length: K381X, and the C-terminus truncation: D373X) using PolyFect Transfection Reagent according to the manufacturer’s protocol (Qiagen) and as previously described^28^. HEK-293 cells expressing MSD1 constructs (~35 kDa) were maintained in DMEM (Wisent) supplemented with non-essential amino acids (Life Technologies) and 10% FBS (Wisent) at 37 °C with 5% CO_2_ (HEPA incubator, Thermo Electron Corporation)^6,9^. These cells were grown at 37 °C (24 h) ± VX-809 (3 μM, Selleck Chemicals), VX-661 (1 μM, Selleck Chemicals) or C18 (6 μM, Cystic Fibrosis Foundation Therapeutics), lysed in modified radioimmunoprecipitation assay buffer (50 mM Tris-HCl, 150 mM NaCl, 1 mM EDTA, pH 7.4, 0.2% (v/v) SDS, and 0.1% (v/v) Triton X-100) containing a protease inhibitor cocktail (Roche) for 10 min, and soluble fractions were analyzed by SDS-PAGE on 4–12% gels. After electrophoresis, proteins were transferred to nitrocellulose membranes and incubated in 5% (w/v) milk, and MSD1 constructs were detected using the human CFTR-MSD1-specific (amino acids 27–34)^29^ murine mAb MM13–4 (1:10000, University of North Carolina Chapel Hill, NC), horseradish peroxidase-conjugated goat anti-mouse IgG secondary antibody (1:5000) and exposure to film for 1–5 min. Calnexin was used as a loading control, and detected using a Calnexin-specific rabbit Ab (1:5000, Sigma-Aldrich), horseradish peroxidase-conjugated goat anti-rabbit IgG secondary antibody (1: 5000) and exposure to film for 1–5 min. Expression levels of MSD1 constructs were quantitated by densitometry using ImageJ software (version 1.42Q, National Institutes of Health) and reported values are normalized to Calnexin expression levels.

### Statistical Analysis

All data are represented as mean ± S.E.M. unless otherwise noted. Non-paired Student’s t-tests, one-way analysis of variance (ANOVA) and Fisher’s exact test were used as appropriate, and *P* values less than 0.05 were considered significant.

## Results

### CFTR Mutation Landscape and Structural Context

Three *CFTR* mutation databases: ABCMdb, CFTR1 and CFTR2, were explored to identify *CFTR* mutations for which the context in the CFTR tertiary structure could be investigated. To this end, only exonic CFTR mutations and their allele frequencies were retrieved from these databases and mapped to their corresponding residue positions in the protein structure. The CFTR structures used for this analysis were a previously described and biochemically validated homology-based model of full-length CFTR in the open-channel state^18^, and a recently reported cryo-EM partial structure of CFTR in the closed-channel state^19^. Boundaries of the CFTR structural domains were defined as follows: MSD1 (1–380), NBD1 (381–630), R-domain (631–849), MSD2 (850–1170), NBD2 (1171–1480), with the numbers under parentheses indicating the first and last residues of the corresponding domain (**Figure S1A and S1B**).

ABCMdb curates the scientific literature and aggregates all reports for a given user-defined search regarding mutations in ATP-Binding Cassette (ABC) proteins, CFTR included. Importantly, this database contains validated disease-causing mutations, unvalidated disease-associated mutations, as well as those used to experimentally investigate CFTR structure and function. For example, data for F508 includes the disease mutation: F508del, as well as all possible amino acid substitutions, and therefore contains the greatest number of ‘counts’ (i.e. twenty) for a single residue position within the CFTR amino acid sequence. Accordingly, this database contains many non-disease-causing mutations. However, such mutations still provide useful insight into the location within the CFTR sequence, which have been biochemically investigated in greater detail.

In total, 1,955 exonic *CFTR* mutations from the ABCMdb were mapped to the CFTR structures (**Figure S2**). Analysis of the structurally mapped mutations reveals that the R-domain, MSD2 and NBD2 are less-well studied compared with MSD1 and NBD1 of CFTR. A somewhat smaller number of mutations were mapped onto the CFTR protein structures from the other two databases: 1,308 mutations from CFTR1 (**Figure 1A and 1B**), and 235 mutations from CFTR2 (**Figure 2A and 2B**). The CFTR1 database represents physician and scientist catalogued mutations, and likely contains the oldest and most comprehensive collection of both validated disease-causing and unvalidated disease-associated *CFTR* mutations. The CFTR2 database contains a smaller collection of *CFTR* mutations, but the corresponding data are enriched for validated disease-causing variants, as they correlate genetic information to biochemical (*in vitro*) and clinical (*in vivo*) parameters and benchmarks (e.g. lung function and pancreatic sufficiency)^7^. Furthermore, the CFTR2 database reports allele frequencies for CF patients mainly in North America and Europe, and thus represents a substantial subset of the worldwide patient population. This dataset is useful for epidemiological studies, and provides a holistic view of the large-scale *CFTR* mutation landscape.

**Figure 1:**
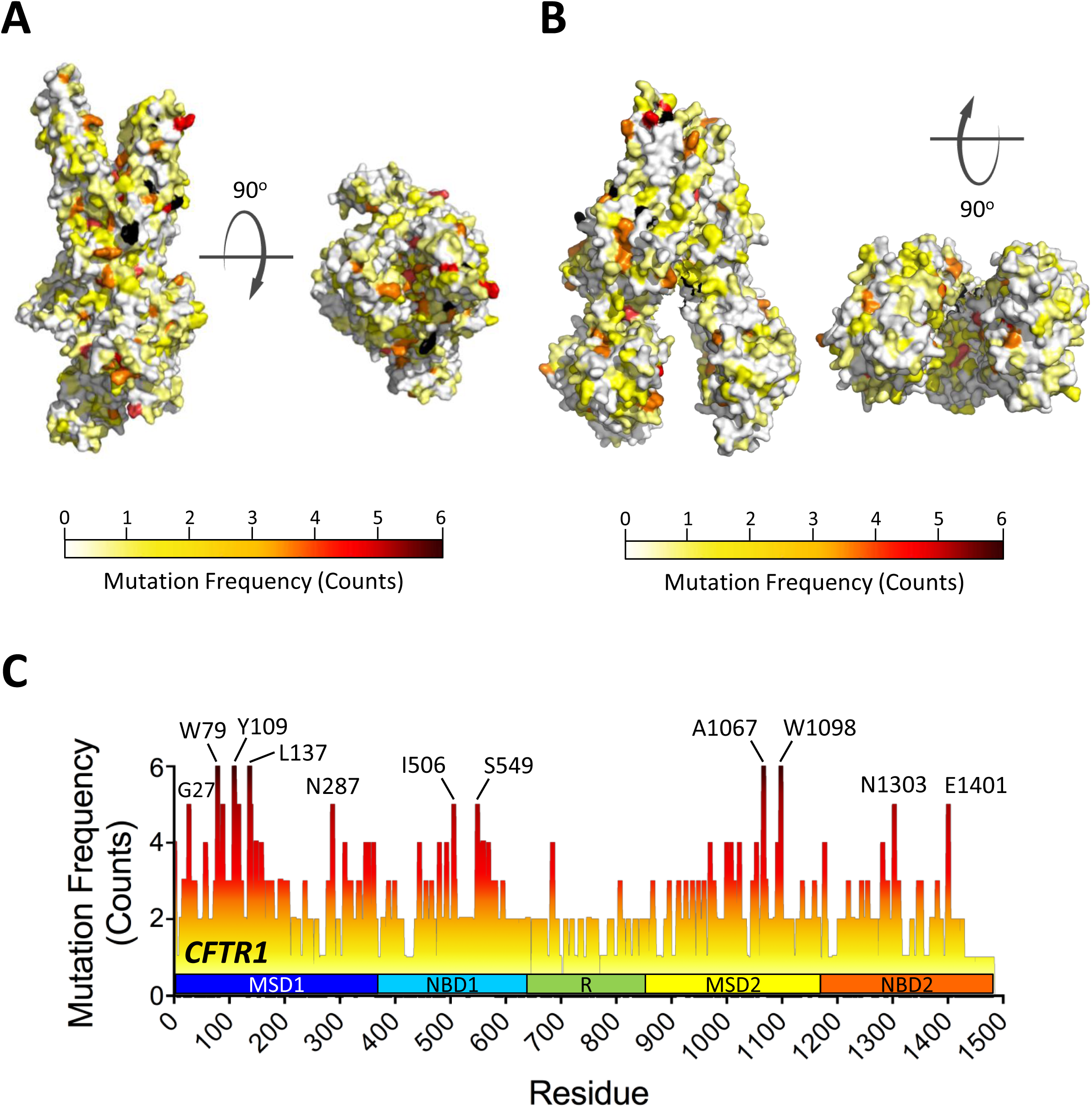
Computational mapping of exonic *CFTR* mutation frequencies from the CFTR1 mutation database to CFTR protein structures. (**A**) Side and top-down views of the full-length CFTR homology model (surface representation) in the open-channel (active) state, and (**B**) side and bottom-up views of the cryo-EM-derived partial structure of CFTR (surface representation) in the closed-channel (inactive) state coloured according to exonic *CFTR* mutation frequencies (counts) reported in the CFTR1 mutation database. (**C**) Histogram of exonic *CFTR* mutation frequencies (counts) as reported in the CFTR1 mutation database for each residue in the protein sequence. Highly mutated residue positions are indicated.

**Figure 2:**
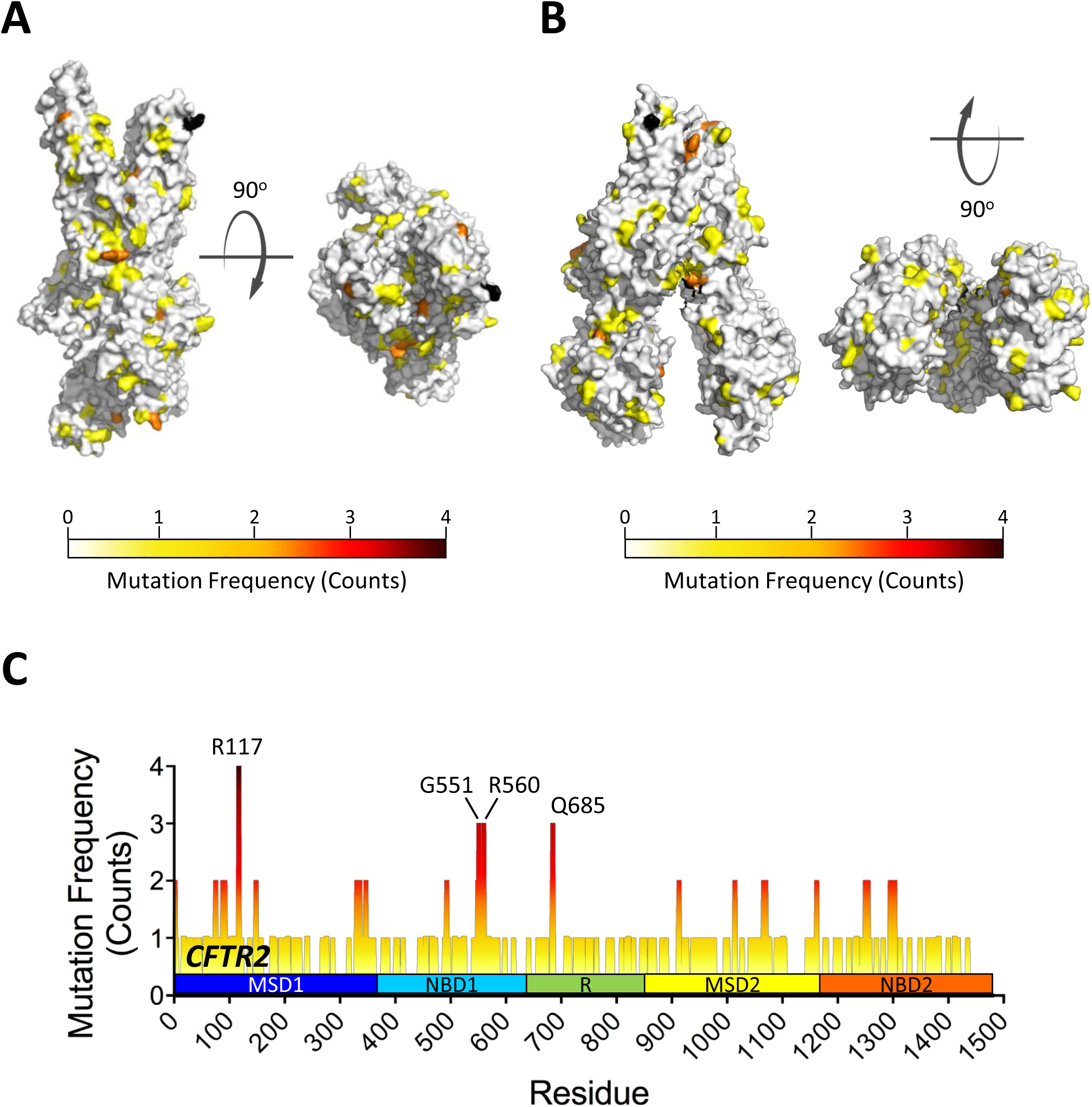
Computational mapping of exonic *CFTR* mutation frequencies from the CFTR2 mutation database to CFTR protein structures. (**A**) Side and top-down views of the full-length CFTR homology model (surface representation) in the open-channel (active) state, and (**B**) side and bottom-up views of the cryo-EM-derived partial structure of CFTR (surface representation) in the closed-channel (inactive) state coloured according to exonic *CFTR* mutation frequencies (counts) reported in the CFTR2 mutation database. (**C**) Histogram of exonic *CFTR* mutation frequencies (counts) as reported in the CFTR2 mutation database for each residue in the protein sequence. Highly mutated residue positions are indicated.

Importantly, the spatial distribution of mapped mutation counts differed between the ABCMdb, CFTR1 and CFTR2 mutation databases. Many mutations from the CFTR1 database mapped to all of the domains (except for the R-domain for which mutations were under-represented) (**Figure 1C**). A smaller collection of mapped mutations was contributed by the CFTR2 database, which had at least one dominant mutation in each of the first three domains (i.e. MSD1, NBD1 and the R-domain) (**Figure 2C**). Overall however, a greater proportion of known mutations from all three databases were located in the first two N-terminal domains of CFTR (MSD1 and NBD1), suggesting that these two domains play an important role in CFTR structure and function.

### Mutant Allele Frequencies and Disease-Causing Mutations Cluster at Distinct CFTR Interdomain Interfaces

In total, 128,354 mutant alleles from the CFTR2 database were mapped onto the tertiary structure of CFTR (**Figure 3A**). Six amino acid positions (F508, G542, G551, R553, W1282, N1303) comprised a majority of the mutant alleles (110,585, ~86% of the total), including 98,735 alleles (~77%) for the F508del mutation alone, and upon further inspection we found that these six positions were spatially co-localized in the tertiary structure of CFTR (**Figure 3B and 3C**). This co-localization (AID of 18.75 Å) was found to be statistically significant (p-value ~0.0003) (**Figure 3D**; see Methods for details). More specifically, these six residues are part of two distinct interdomain interfaces (NBD1:NBD2 and NBD1:ICL4) which have been previously described as being integral to CFTR structure and function, and accordingly, represents a region of mutation clustering that is over-represented in the CF patient population when compared to all of the alleles reported in the CFTR2 database^30–33^.

**Figure 3:**
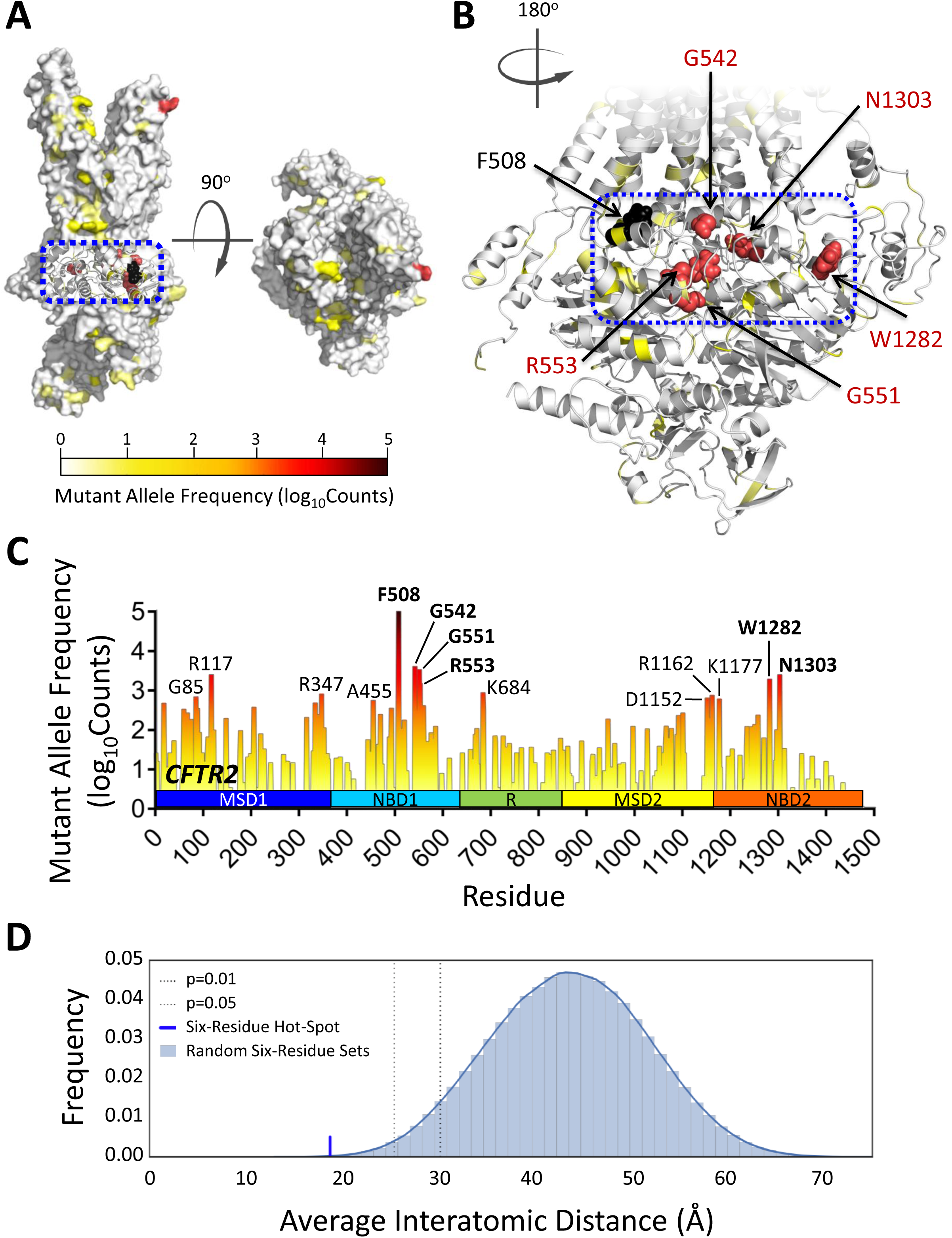
Computational mapping of exonic *CFTR* mutant allele frequencies from the CFTR2 mutation database to a structural model of CFTR. (**A**) Side and top-down views of the full-length CFTR homology model (surface representation) in the open-channel (active) state coloured according to exonic *CFTR* mutant allele frequencies (log_10_counts) reported in the CFTR2 mutation database. The boxed (blue) region of CFTR indicates a mutation cluster. (**B**) Close-up view of mutation cluster. Six residues with top-ranking mutant allele frequencies (log_10_counts) are labelled, and include: F508, G542, G551, R553, W1282 and N1303. (**C**) Histogram of exonic *CFTR* mutant allele frequencies (log_10_counts) as reported in the CFTR2 mutation database for each residue in the protein sequence. Highly mutated residue positions are indicated. (**D**) Distribution of AID values for randomly selected six-residue sets relative to the AID of the six-residue cluster (F508, G542, G551, R553, W1282, N1303).

To assess the frequencies of disease-causing mutations at residue positions of the NBD1:NBD2 and NBD1:ICL4 interdomain interfaces, we first identified amino acids that contribute to these interfaces (see **Figure S3** and Methods for details). This revealed that the NBD1:NBD2 interface was more extensive (94 residues) than the NBD1:ICL4 interface (29 residues) as illustrated in **Figure 4 and Table 1**. We then computed the fraction of residues (i.e. ratio) with disease-causing mutations for the full-length CFTR protein to be 0.88 (i.e. 1,308 exonic *CFTR* mutations among 1,480 total residues, as reported in the CFTR1 database). The corresponding fractions for the NBD1:NBD2 and NBD1:ICL4 interfaces were 1.06 (i.e. 88 mutations among 94 residues, p>0.05) and 1.60 (i.e. 41 mutations among 29 residues, p<0.05), respectively (**Table 1**). These data indicate that these two interdomain interfaces have, on average, a larger fraction of mutated residues than full-length CFTR. Importantly, NBD1:ICL4 was found to harbour a statistically significant cluster of mutations, in agreement with the current paradigm that this region plays a crucial role for competent CFTR biosynthesis, stability and activity, as demonstrated by the major mutation (F508del) which is located at this interface^32,33^.

**Figure 4:**
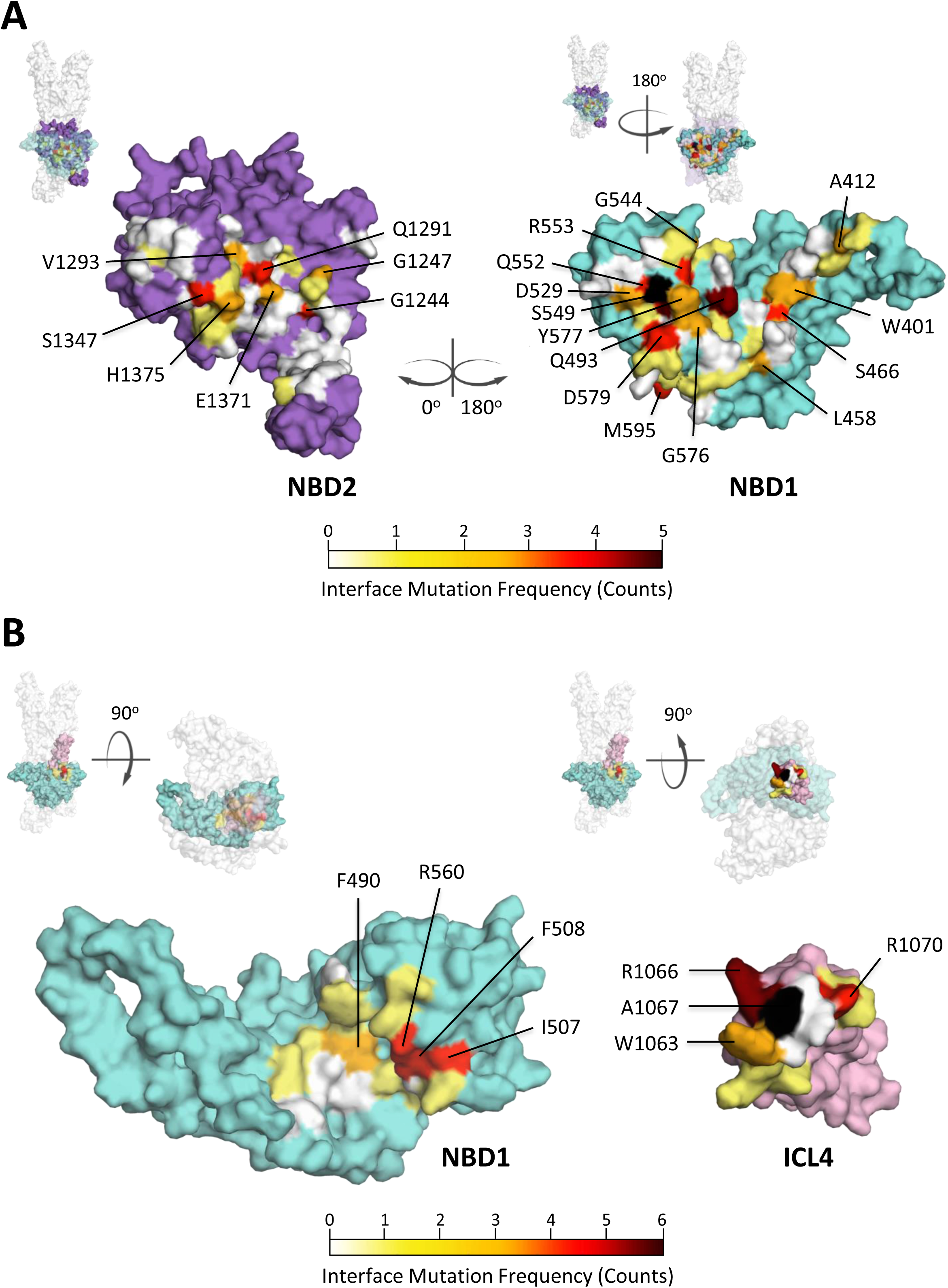
Identification of exonic *CFTR* mutations at NBD1:NBD2 and NBD1:ICL4 interdomain interfaces in the structural model of CFTR. (**A**) NBD1:NBD2 and (**B**) NBD1:ICL4 interdomain interfaces in the full-length CFTR homology model (surface representation) the open-channel (active) state. Interface residues (i.e. those < 4 Å from the opposing domain) are coloured white, and exonic *CFTR* mutations (counts) found within interface residues are coloured according to values reported in the CFTR1 mutation database. Cyan, NBD1 (residues 381–630); purple, NBD2 (1171–1480); pink, ICL4 (1039–1093).

### Prediction of Corrector-sensitive CFTR Residues

The recently determined Cryo-EM structure of CFTR represents an unprecedented opportunity for identifying the binding sites of corrector ligands, as this structure represents the closed-channel conformation of the protein to which such ligands are believed to bind. Identifying the long sought binding site, or sites, of such ligands should help gain insight into the mechanism-of-action of these molecules and allow for patient stratification based on their mutation-sensitive ‘theratype’.

To this end, we consider the Class I corrector ligands VX-809, VX-661 and C18, which share a common scaffold (pharmacophore), but also display some differences (**Figure S4A**). Previous studies have shown that VX-809 stabilizes the MSD1 domain of CFTR^27,28,34^. This stabilizing effect was abolished in MSD1 constructs that lacked the C-terminal segment comprising residues 374–380^27^. In addition, the deletion of only residues 371–375 from full-length CFTR produced a severe folding defect that could not be corrected by VX-809^27^. It therefore seems of interest to explore this region of CFTR for corrector binding sites. This choice was further confirmed by immunoblotting experiments carried out in this study. These experiments show that deletion of residues 373–380 abrogates corrector-mediated increase in MSD1 abundance, a common measure of the presence of folded protein, for all three Class I correctors (VX-809, VX-661 and C18) (**Figure 5A and 5B**). This observation suggests that all three ligands interacted with the C-terminal region of MSD1, in agreement with previous findings on VX-809 alone.

**Figure 5:**
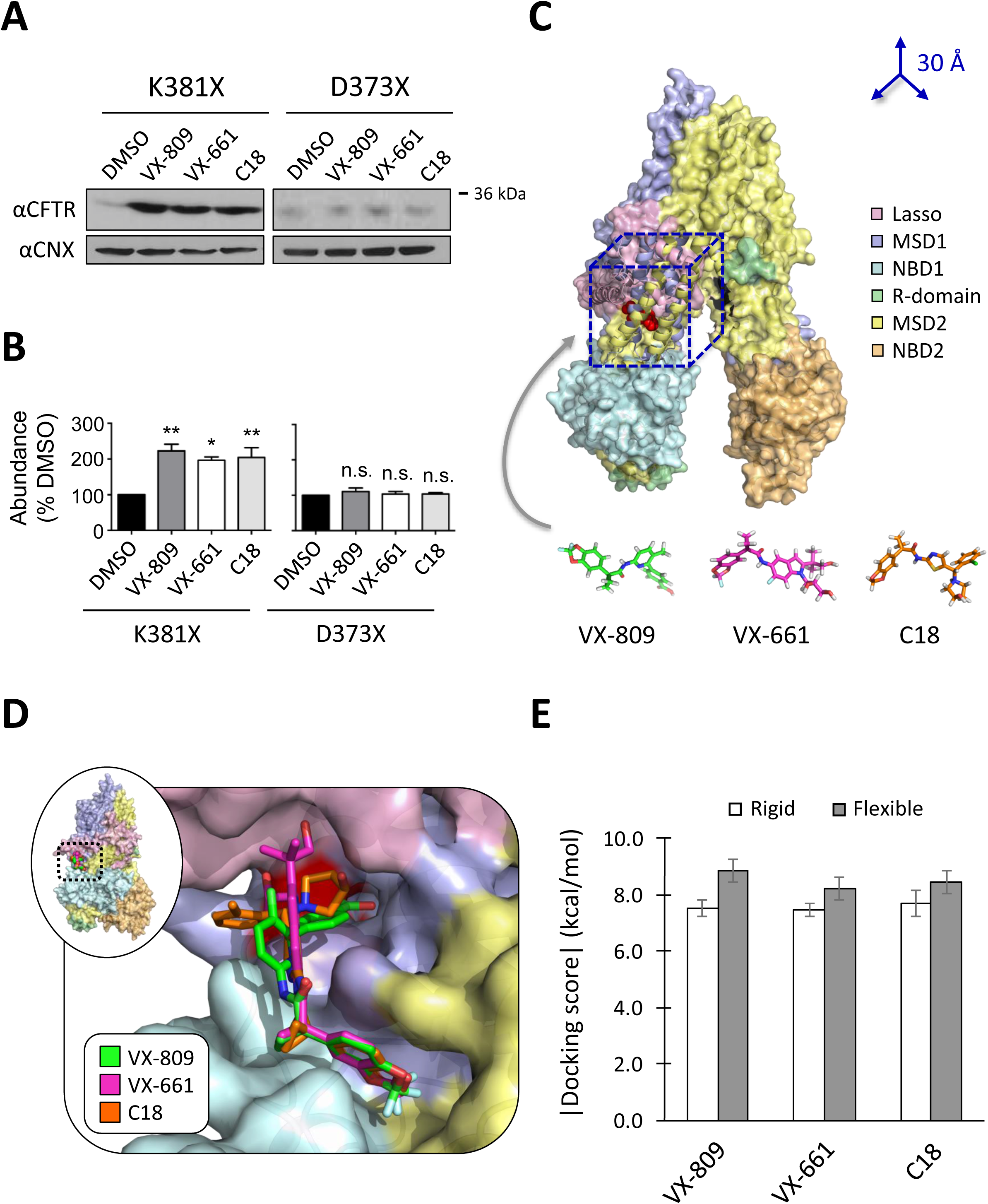
Identification of the putative Class I corrector binding pocket using molecular docking. (**A**) Immunoblots and (**B**) quantitative analysis of HEK-293 cells transiently expressing CFTR MSD1 fragments (i.e. full-length: K381X, and the C-terminus truncation: D373X) following chronic treatment (24 h) with correctors VX-809 (3 μM), VX-661 (1 μM) and C18 (6 μM) (n=3). Calnexin was used as a loading control. **, p<0.01; *, p<0.05; n.s., not significant. (**C**) The cryo-EM-derived partial/core-structure of CFTR in the closed-channel (inactive) state served as a template for molecular docking studies. The 30×30×30-Å cube representing the docking boundary, localized around residues F374-L375 (red spheres), is shown (dashed, dark blue lines); CFTR is shown in surface representation and coloured by domain as described in the legend. (**D**) Close-up view of binding modes for VX-809 (green), VX-661 (magenta) and C18 (orange), demonstrating alignment of the common scaffold, following molecular docking into the top-ranking and putative corrector binding site; CFTR is shown in surface representation and domains are coloured as in panel **C**. (**E**) Mean docking scores for top-scoring binding modes of Class I correctors following rigid (white bars) or flexible (gray bars) docking to the cryo-EM-derived structure of CFTR in the closed-channel state (n=9).

Docking calculations for all three ligands were hence carried out in the vicinity of residues F374 and L375 using the bounded volume described in the Methods and illustrated in **Figure 5C**. Assuming a similar mode of action for the three ligands, we favored docking poses in which the common scaffold of these molecules was closely aligned. The docking results, consisting of a top-scoring binding pocket, and the associated corrector conformations are shown in **Figures 5D and 5E**. All three modeled CFTR-ligand complexes featured comparable docking scores and involve very similar interactions with CFTR residues lining the pocket (**Table S1**). The majority of the residues making contact with the ligands are from MSD1 (i.e. lasso motif, ICL1 and C-term), a few are from the N-terminus of NBD1, and only a couple of residues at most from MSD2 (i.e. ICL4). The side-chain of one residue in particular, K166 (ICL1), appears to play an important role in corrector recognition. It was positioned closest (~4.0 Å) to the plane of the common scaffold in the top-scoring CFTR-corrector complexes, allowing the ε-amino group of the lysine side-chain to possibly coordinate with the aromatic ring or carbonyl oxygen of this scaffold of each corrector so as to form cation-π interactions or H-bonds, respectively (**Figure S4B**). It is noteworthy that the ligands make no contact with the side-chain of F374 and minimal contact with the side-chain of L375, consistent with the finding that mutations of these residues to Alanine has little effect on the potency of VX-809^27^.

## Discussion

In this study, the known repertoire of exonic mutations and mutant allele frequencies in *CFTR*, compiled in three public databases, was mapped onto the three-dimensional structural models of the CFTR protein. Analysis of the spatial distribution of the mapped mutations revealed that disease-causing mutations tend to occur more frequently in certain regions of the CFTR structure, which likely play a strategic role in conferring the thermodynamic and dynamic properties required for protein function.

Mapping *CFTR* mutations from the ABCMdb, which includes validated disease-causing mutations, unvalidated disease-associated mutations, and those of experimental origin, showed that the N-terminal domains of CFTR (MSD1 and NBD1) have been more extensively probed by mutation studies. The greater focus on the N-terminal domains may be due to the occurrence in these domains of two relatively more common disease-causing mutations, R117H and F508del, and perhaps also to the increased influence that residues in the N-terminal half of this large polypeptide may have on proper folding of the full CFTR protein.

In contrast, mapping disease-causing mutations only from the CFTR1 and CFTR2 databases to the CFTR protein structures revealed that, for CFTR1, such mutations are found in all domains across the structure, while for CFTR2, where the mutations data is more limited but enriched for validated variants, MSD1 and NBD1 contain the greatest abundance of mutated residues. Further, analysis of the spatial distribution of amino acid positions of the most frequent mutant alleles derived from the CFTR2 database, revealed a statistically significant cluster in the tertiary protein structure containing six of the most common disease-causing mutations; this spatial distribution was not completely apparent in the primary sequence. We speculate that selection pressures (e.g. environmental stresses) played a role in giving rise to this cluster, possibly supporting the *heterozygote advantage* hypothesis^35–38^. The latter suggests that carriers (i.e. heterozygotes) of a mutant *CFTR* allele are thought to have had a survival advantage compared to individuals with wild-type, and thus pressure-susceptible alleles; the leading hypotheses, although controversial among CF researchers, propose a survival advantage against cholera, typhoid fever and/or tuberculosis^35–38^.

In addition, we found that disease-causing mutations were clustered at two distinct interdomain interfaces, NBD1:NBD2 and NBD1:ICL4, with the cluster at the latter interface being statistically significant (p<0.05). This observation confirms that these regions are fragile ‘linchpins’, essential for intramolecular communication and facilitating conformational changes required for CFTR function, and further, that mutation of individual residues within these interfaces is sufficient to destabilize and/or hamper proper tertiary assembly of functionally competent states of the protein^29,30^. This is in agreement with previous studies, which showed that residues buried within domain-domain interfaces are commonly associated with disease-causing mutations of many proteins in the human proteome^39^.

Two models for the three-dimensional structure of the CFTR protein were used for the mapping analysis, the full-length CFTR homology model representing the open-channel, active state of the protein and the cryo-EM partial structure of CFTR in the closed-channel, inactive state. Neither model is of high accuracy by current standards of protein crystal structures. But this may only marginally affect the observations made from the mutation mapping analysis, as the latter should not critically depend on model accuracy. On the other hand, the cryo-EM structure has the advantage of being the first experimentally-derived CFTR structure with amino acid side-chain resolution. The asset of the homology model is that it is truly full-length. It includes built atomic coordinates for the structurally flexible R-domain (residues 631–849), which contains approximately 10% of all *CFTR* mutations. In addition, the model for the MSD-NBD dimer portion is based on the canonical scaffold, which is well-represented in the structurally-characterized Protein Data Bank (PDB). This model represents the ATP-bound state of CFTR, which correlates with a large set of structural homologues in the PDB (i.e. ATP-bound, non-mammalian ABC transporters) and is in agreement with several biochemical studies, further supporting its validity^18^.

In the last part of our study, we investigated the putative binding site of three Class I correctors molecules, VX-809, VX-661 and C18. We confirmed experimentally that all three ligands stabilized MSD1 of CFTR, requiring the C-terminal segment of this domain to elicit their corrector action, as reported for VX-809 in other studies^27,28,34^. The three ligands were then computationally docked onto a 30-Å**^3^** region centered at the C-terminal region of MSD1 (i.e. residues F374-L375) in the cryo-EM structure of CFTR, and poses which demonstrated alignment of the pharmacophore (i.e. common scaffold) of these three small molecules suggested the most plausible binding pocket. This enabled the identification of a sizable binding pocket with least 12 residues of the CFTR protein (a sufficient number for ligands of these molecular weights: i.e. ~450–520 Da), many of which were in MSD1, engaging in interactions with the bound ligands (**Table S1**). Our findings that the identified corrector binding pocket comprises residues from different domains (i.e. lasso motif, ICL1 and C-term of MSD1, N-terminus of NBD1, and ICL4 of MSD2; **Table S1**) further elaborate on previous *in silico* docking studies, which propose that a composite, multi-domain pocket comprised of residues within the NBD1:ICL4 interface (near the major mutation: F508del) is likely to accommodate VX-809^24,25^.

More specifically, we found that the three non-pharmacophore hydroxyl groups of VX-661, which are not featured in the investigational and first-generation corrector molecules: C18 and VX-809, respectively (**Figure S4A**), formed H-bonds with residues E54, W57 and K163 in the predicted pocket (**Figure S4B**), possibly explaining the *in vitro* effectiveness of VX-661 over these other correctors. Furthermore, binding poses for all three correctors position the pharmacophore in a groove comprising residues K166, Y380, T382, R1066 and Q1071, and interacting more closely with K166. In the complex with VX-661, the positively charged epsilon amino group of K166 was appropriately positioned towards the aromatic benzodioxole group to form a potential cation-π interaction^40–42^. C18 featured a slight rotation of the K166 side-chain that places its charged group in an appropriate geometry for an H-bond with the corrector’s carbonyl oxygen of the pharmacophore (data not shown). These different options for the K166 side-chain to form stabilizing interactions with different ligands or possibly with the same ligand, support the predicted involvement of this residue in corrector binding. Furthermore, our future studies will investigate the role of K166 in coordination and subsequent binding of these Class I correctors using a site-directed mutagenesis approach.

Finally, it is noteworthy that disease-causing mutations have been reported for several residues in the predicted binding pocket (e.g. R1066C/S/H/L; see **Table S1**). It is therefore possible that the Class I correctors analyzed here may repair structural defects caused by these variants (locally in this pocket and globally throughout CFTR protein), and potentially become therapeutically relevant for CF patients bearing these mutations. Taken together, these data suggest that the current CF co-therapy (i.e. Orkambi®, but mainly lumacaftor) may benefit this patient subpopulation. Furthermore, it is clear that additional investigations using mutational and biophysical analyses are needed to confirm the predicted binding pocket and the proposed contributions of various CFTR residues therein to the binding of the three Class I correctors.

## Table Captions

**Table 1: Clustering of exonic *CFTR* mutations at the NBD1:ICL4 interdomain interface of CFTR.** The CFTR1 mutation database was used to map exonic *CFTR* mutation frequencies (counts) onto interdomain interface residues of the full-length CFTR homology model in the open-channel (active) state using a proximity-measure (i.e. residues < 4 Å from an opposing surface or ‘face’ were deemed interdomain positions). Relative mutation frequencies at interdomain interfaces (i.e. NBD1:NBD2 and NBD1:ICL4) were compared to the relative mutation frequency of full-length CFTR to assess significance. *, p<0.05.

## Acknowledgments

### Author contributions

The overall design of the study was by S.V.M.; S.V.M., V.M.S., A.S.S., S.S.M., G.W., M.L., O.L., L.D.M., C.E.B. and A.W. performed experiments, analyzed and/or interpreted results. The manuscript was primarily written by S.V.M. with input from all authors.

### Funding

This research did not receive any specific grant from funding agencies in the public, commercial, or not-for-profit sectors.

### Competing interests

All authors affiliated with Cyclica are employees or advisors to the company. All other authors declare no conflicts of interest.

## Supplementary Figure Legends

**Figure S1:**
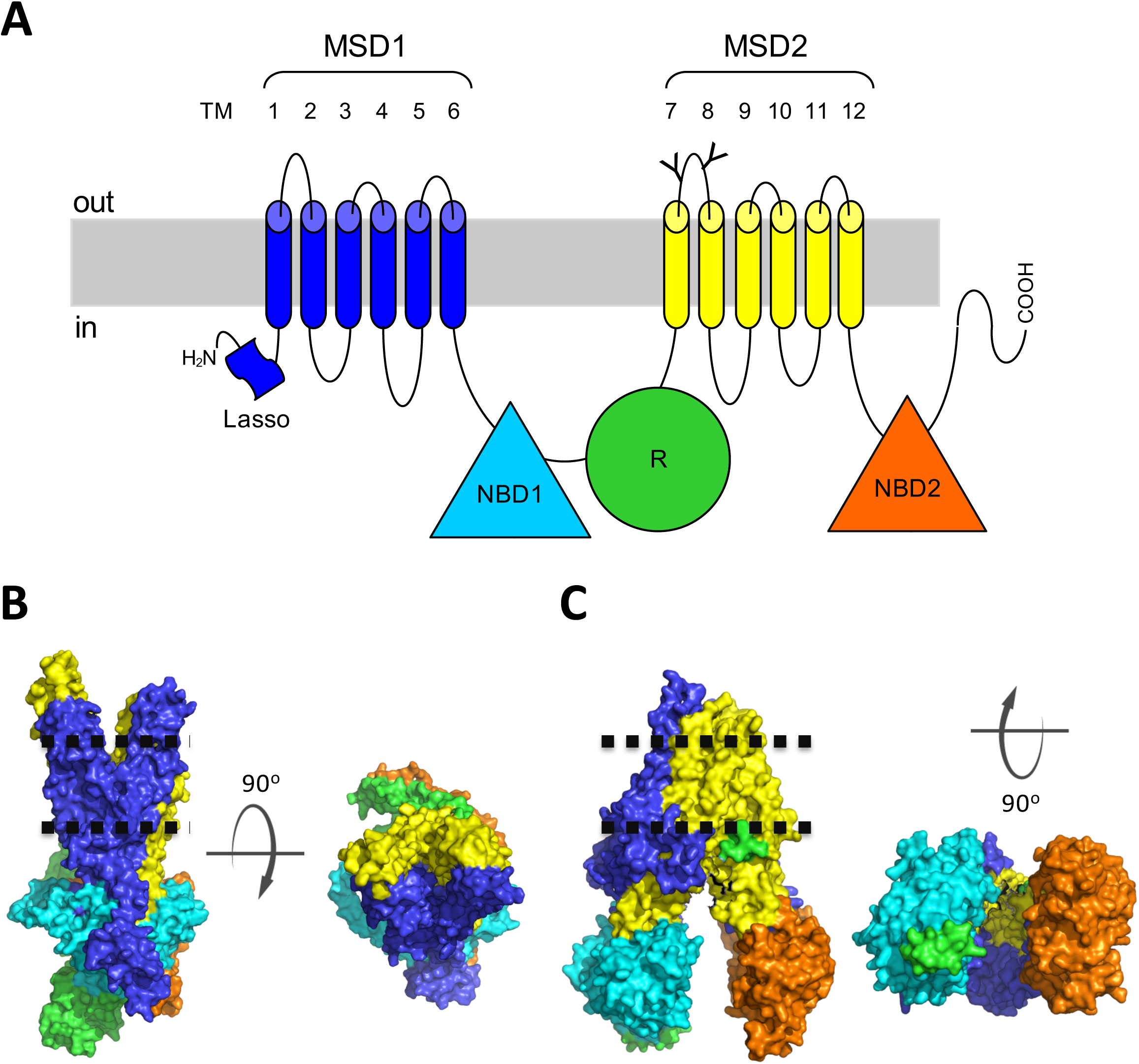
Topological model and domain-coloured tertiary structures of CFTR. (**A**) A topology model of CFTR showing MSD1 (blue, residues 1–380), NBD1 (cyan, residues 381–630), R-domain (green, residues 631–849), MSD2 (yellow, residues 850–1170) and NBD2 (orange, residues 1171–1480). The ‘Y-shaped’ symbols represent N-glycosylation sites. TM, transmembrane α-helix. The relative position of the plasma membrane is shown in gray. (**B**) Side and top-down views of the full-length CFTR homology model (surface representation) in the open-channel (active) state, shown in surface representation. (**C**) Side and bottom-up views of the cryo-EM-derived structure of CFTR (lacking a majority of the R-domain, i.e. residues 646–843) in the closed-channel (inactive) state, shown in surface representation. Domains are coloured as in panel **A**, and the relative position of the plasma membrane is shown with a dotted line.

**Figure S2:**
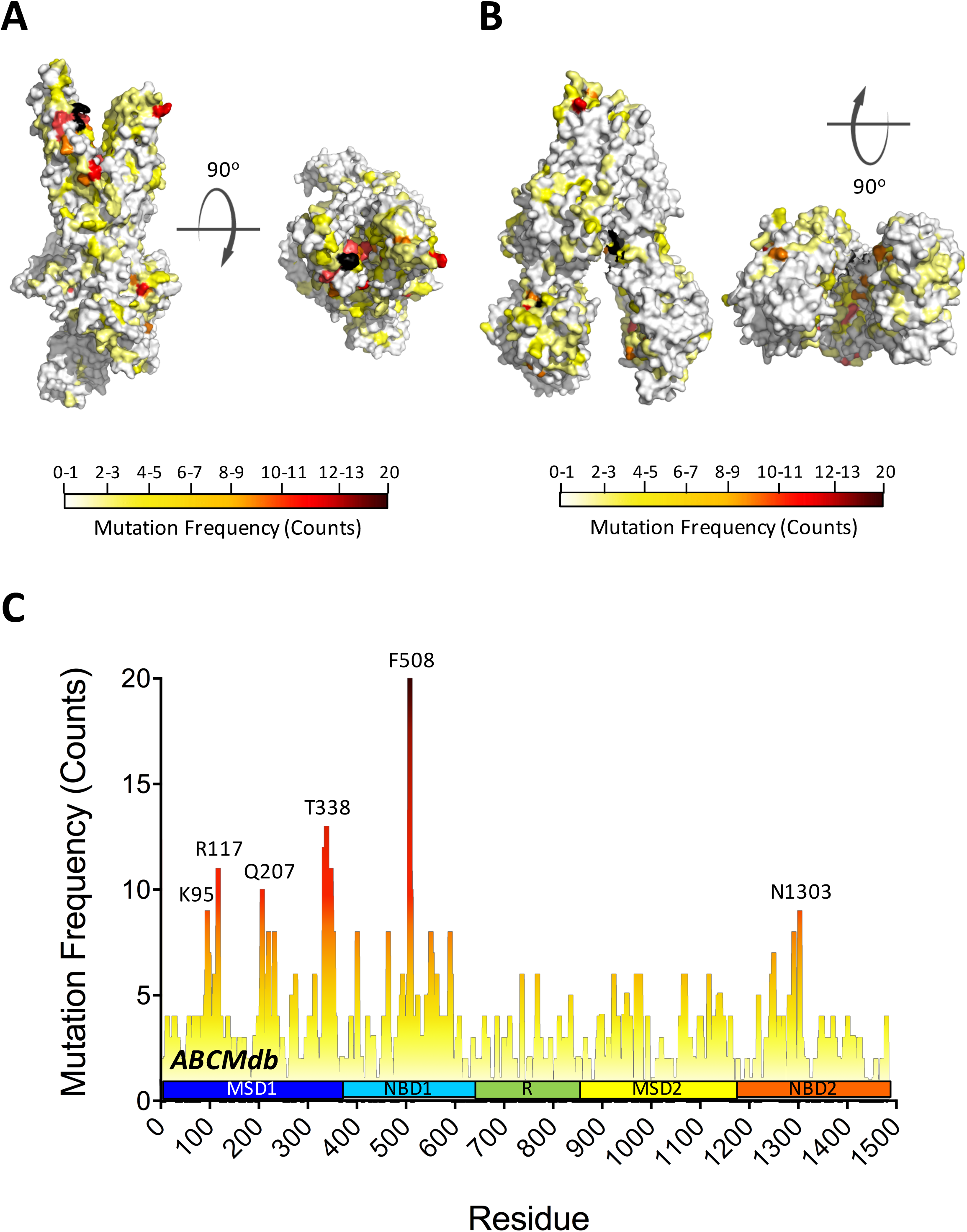
Computational mapping of exonic *CFTR* mutation frequencies from the ABC mutation database to CFTR protein structures. (**A**) Side and top-down views of the full-length CFTR homology model (surface representation) in the open-channel (active) state, and (**B**) side and bottom-up views of the cryo-EM-derived partial structure of CFTR (surface representation) in the closed-channel (inactive) state coloured according to exonic *CFTR* mutation frequencies (counts) reported in the ABC mutation database. (**C**) Histogram of exonic *CFTR* mutation frequencies (counts) as reported in the ABC mutation database for each residue in the protein sequence. Highly mutated residue positions are indicated.

**Figure S3:**
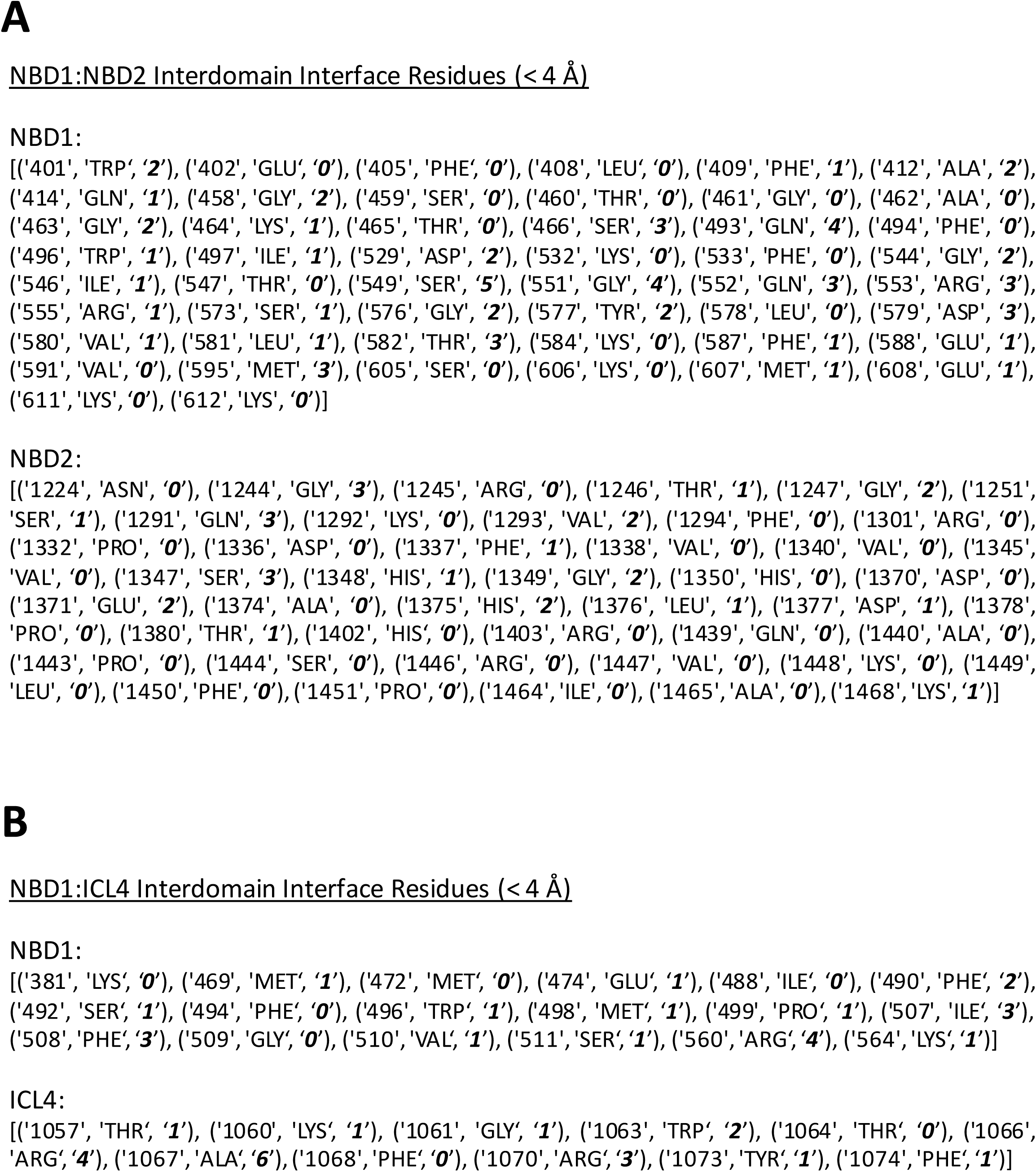
List of CFTR residues located at NBD1:NBD2 and NBD1:ICL4 interdomain interfaces. (**A**) NBD1 and NBD2 residues found within 4 Å of the opposing face of the NBD1:NBD2 interdomain interface, as generated from a PyMOL-based proximity-measure script. Syntax includes ‘X’, ‘Y’ and ‘Z’ coordinates as: position, native residue, and number of mutations (in bold italics), respectively. (**B**) NBD1 and ICL4 residues found within 4 Å of the opposing face of the NBD1:ICL4 interdomain interface, as generated from a PyMOL-based proximity-measure script. Syntax includes ‘X’, ‘Y’ and ‘Z’ coordinates as: position, native residue, and number of mutations (in bold italics), respectively.

**Figure S4:**
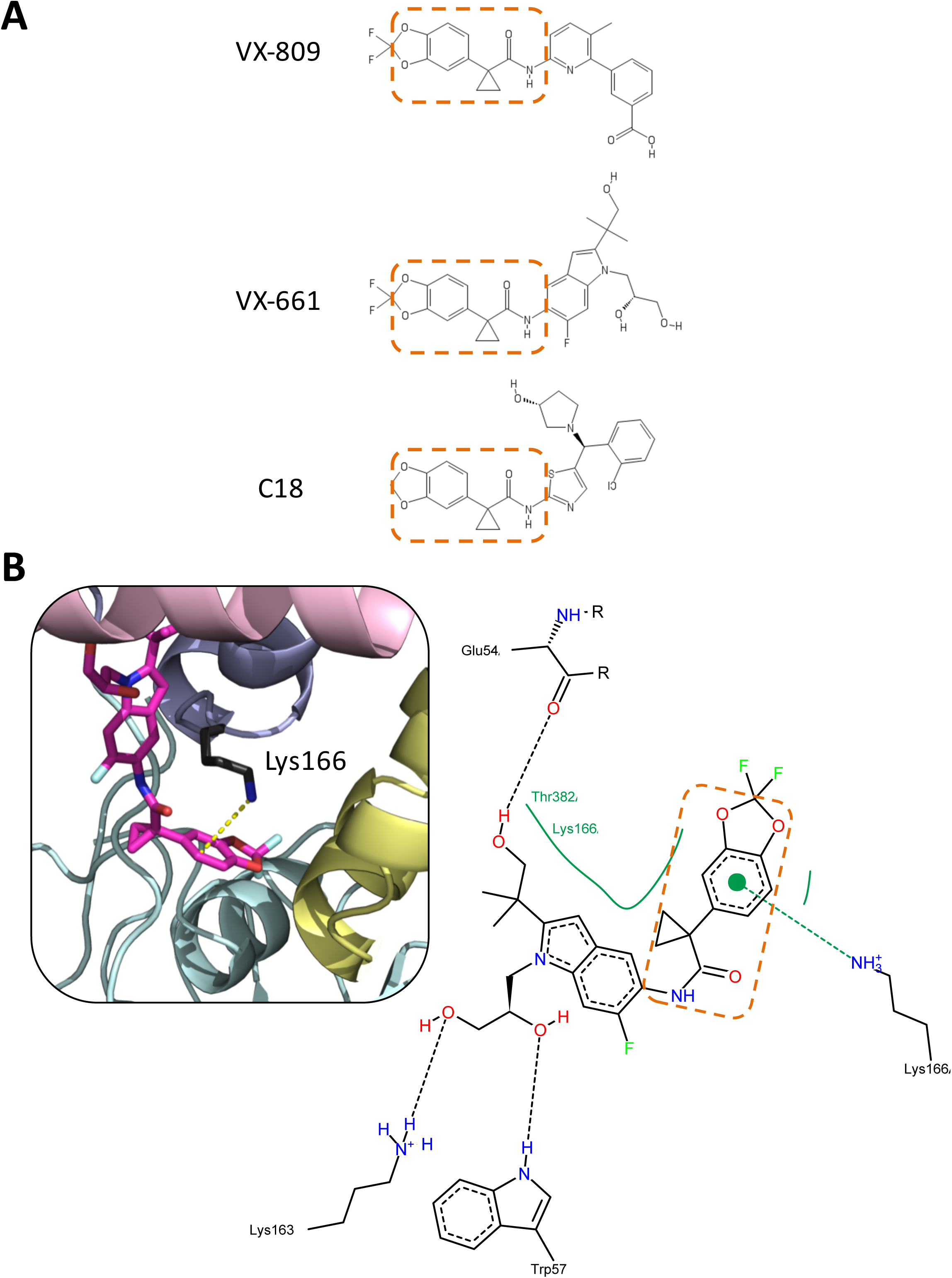
Chemical structures of VX-809, VX-661, C18, and the predicted binding mode of VX-661 in the putative corrector binding pocket. (**A**) Chemical structures of each Class I corrector. The common scaffold or ‘pharmacophore’ of each small molecule is highlighted with a dashed box (orange). (**B**) As an example, the predicted binding mode of VX-661, captured in the putative corrector binding pocket of the cryo-EM-derived partial structure of CFTR in the closed-channel (inactive) state, is shown in the inset (pink, lasso motif; yellow, ICL4 of MSD2; cyan, NBD1; light blue, ICL1 of MSD1; dark gray, K166; magenta, VX-661; dashed yellow line, predicted cation-π interaction). A two-dimensional representation of the predicted binding mode of the example Class I corrector, VX-661, in the putative corrector binding pocket (corresponding to the inset) is also shown, as generated using the PoseView tool from the ZBH Proteins*Plus* webserver (proteinsplus.zbh.uni-hamburg.de).

## Supplementary Table Captions

**Table S1:**
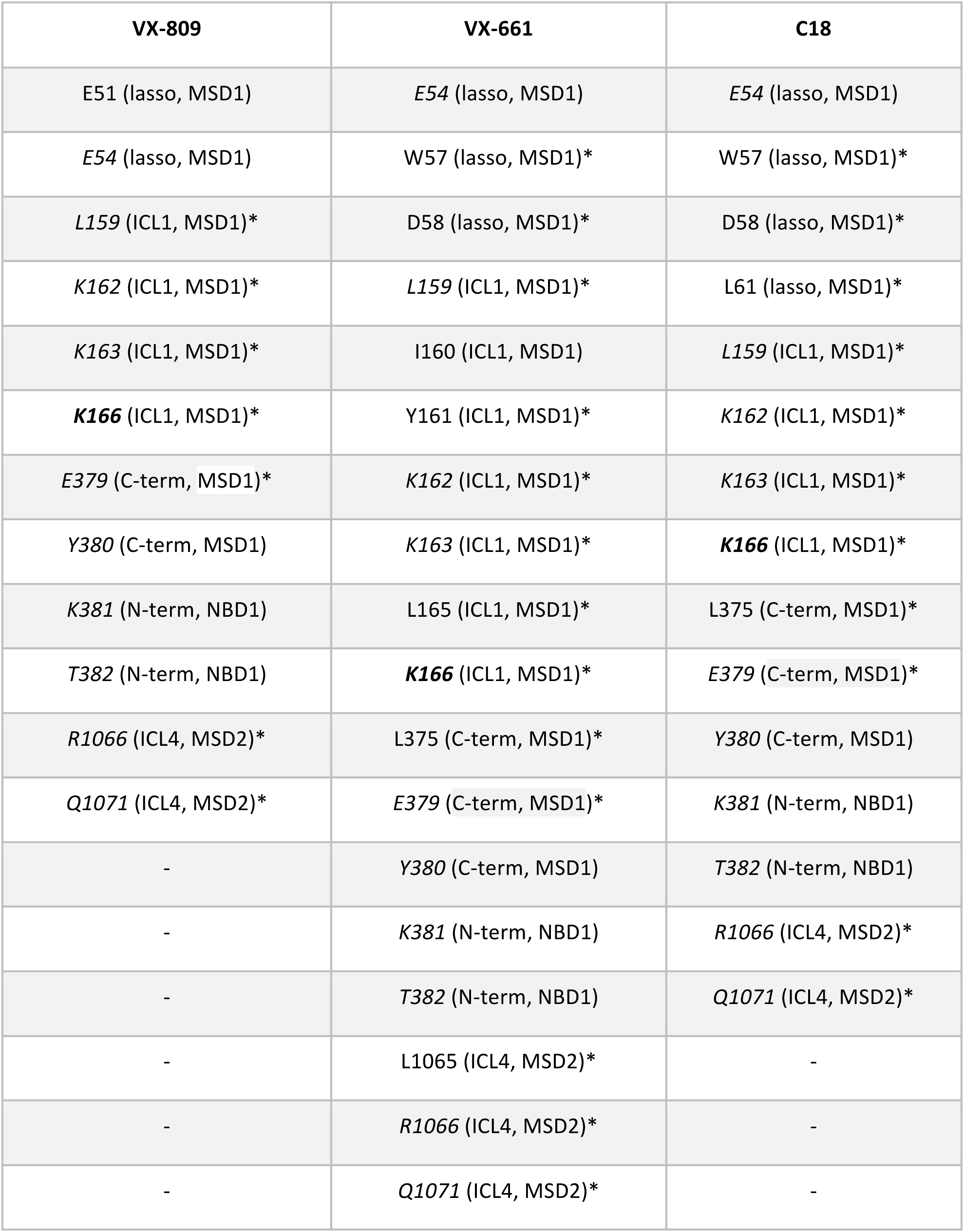
CFTR residues predicted to interact with Class I correctors VX-809, VX-661 and C18. List of residues (subdomain, domain) predicted to be within 5 Å of each respective small molecule corrector following molecular docking studies. Residues in italics are common among the three compounds, and the residue in bold italics (i.e. K166) is found ~4.0 Å of the common scaffold or ‘pharmacophore’ of each corrector. Residues bearing disease-associated missense mutations, as reported in the CFTR1 database, are labelled with an asterisk.

